# Local GPCR density tips the balance of μ-opioid receptor trafficking

**DOI:** 10.64898/2026.02.26.708286

**Authors:** Michael D. Holsey, Alexey Bondar, Peter Geggier, George V. Dukas, Chase M. Webb, Alekhya Govindaraju, Signe Mathiasen, Meritxell Canals, Nevin A. Lambert, Wesley B. Asher, Jonathan A. Javitch

## Abstract

The extent to which local GPCR surface density governs engagement of downstream signaling and trafficking pathways remains unclear. Using single-particle tracking of the μ-opioid receptor (MOR), we show that receptor density differentially regulates G protein signaling and GRK2/3–β-arrestin–dependent receptor trafficking. At low surface density, MORs activate G proteins but fail to enter clathrin-coated structures despite the presence of endogenous GRK2/3 and β-arrestin. Increasing MOR density, co-expressing other class A GPCRs, or elevating GRK2 or β-arrestin abundance rescues agonist-induced MOR trafficking. In contrast, the class B GPCR V2R blocks MOR trafficking at both low and high MOR densities. These results support a model in which increasing class A GPCR density, despite worsening effector-to-receptor stoichiometry, promotes trafficking by forming an affinity matrix that enables reversible GRK2/3 and β-arrestin interactions to be productively used by neighboring receptors in a density-dependent manner, whereas class B GPCRs sequester β-arrestin and block trafficking.

## INTRODUCTION

G protein-coupled receptors (GPCRs) constitute the largest family of cell-surface membrane proteins and are among the most successful targets for therapeutic development; they regulate diverse physiological processes and contribute to numerous human diseases.^1–3^ Upon agonist binding, GPCRs activate heterotrimeric G proteins to initiate downstream signaling, while also engaging GPCR kinases (GRKs) and β-arrestins that limit further G protein coupling and promote receptor trafficking to clathrin-coated structures (CCSs) and internalization.^4–6^ β-arrestins can also scaffold distinct signaling pathways downstream of activated receptors.^7^

An often overlooked but critical factor in these signaling and trafficking processes is the local density of GPCRs at the cell surface, which can be shaped not only by changes in receptor expression and internalization, but also by local membrane organization and interacting proteins. Receptor density can be dynamically regulated and directly influences the potency and magnitude of G protein–mediated signaling in response to ligand binding, consistent with the classical concept of receptor reserve, but how receptor abundance shapes engagement with GRKs, β-arrestins, and trafficking machinery remains poorly understood.^8–12^ The µ-opioid receptor (MOR) is both a prototypical class A GPCR and a critical therapeutic target,^13^ making it an ideal model for understanding how receptor density influences GPCR function. MORs are the primary targets of potent analgesics (e.g., morphine, fentanyl), yet chronic opioid use leads to tolerance, a phenomenon linked to receptor desensitization and β-arrestin-mediated trafficking, ^14–16^ including enhanced receptor phosphorylation, β-arrestin recruitment, and subsequent internalization into clathrin-coated structures.^15,17,18^ MOR surface density fluctuates during tolerance development and varies across different neuronal populations,^15,19–22^ suggesting that local receptor density may be a key determinant of how efficiently MORs engage trafficking pathways and downstream signaling outcomes.

Here, we use single-particle tracking (SPT) to investigate how different cell-surface densities of MOR affect its plasma membrane organization and trafficking to CCSs. SPT enables direct visualization of individual receptor molecules, allowing us to quantify receptor motion dynamics associated with interactions with endocytic structures. Unexpectedly, at very low surface densities, MORs exhibited little to no agonist-induced trafficking to CCSs, despite retaining G protein signaling capacity and the presence of endogenous cytoplasmic effectors, most notably β-arrestins as well as GRK2 and GRK3 (GRK2/3), which are expected to be in stoichiometric excess relative to receptor. These observations indicate that effector abundance alone is insufficient to support MOR trafficking. However, increasing MOR density or co-expressing other GPCRs in the class A β-arrestin-interacting family enabled agonist-induced trafficking of MORs to CCSs, despite a substantially reduced receptor-to-effector stoichiometric ratio. We conclude that receptor density is a critical determinant of MOR engagement with CCSs, revealing a mechanism in which the local membrane density of activated class A GPCRs, together with shared cytosolic effectors, governs trafficking outcomes.

## RESULTS

### MORs at low surface densities show limited interactions with CCSs

To detect individual MORs in the plasma membrane for SPT, we used a Chinese hamster ovary (CHO) cell line that stably expresses amino-terminally SNAPfast (SNAP_f_)-tagged MORs under tetracycline-regulated expression.^23^ This cell line and its integrated receptor-encoding plasmid were engineered to achieve low receptor expression without tetracycline induction while producing substantially higher receptor expression when induced. SNAP_f_-MOR was labeled with the photostable, self-healing organic fluorophore Lumidyne 555p (LD555p)^23–25^ and imaged using total internal reflection fluorescence (TIRF) microscopy (**Methods**) (**Figures 1A and 1B**). In the absence of tetracycline induction, the engineered CHO cells exhibited low surface receptor densities, averaging 0.13 ± 0.04 particles/µm^2^, a density compatible with SPT (**Figures 1C and S1A**).^23^ Individual MOR trajectories in the plasma membrane were generated using the u-track particle tracking algorithm and analyzed with divide-and-conquer moment scaling spectrum (DC-MSS) analysis^26^ to classify receptor motion states. DC-MSS classifies trajectory segments into distinct motion states: free or directed motion, which are mobile diffusion states covering larger membrane areas, and confined motion and immobility, which represent restricted motion states within particular membrane regions^27^ (**Figure 1D**). In the absence of tetracycline induction and agonist stimulation, MORs spent an average of 55% of their total trajectory time in the free diffusion state, 20% in the confined state, and 24% in the immobile state (**Figure 1E**).

**Figure 1.**
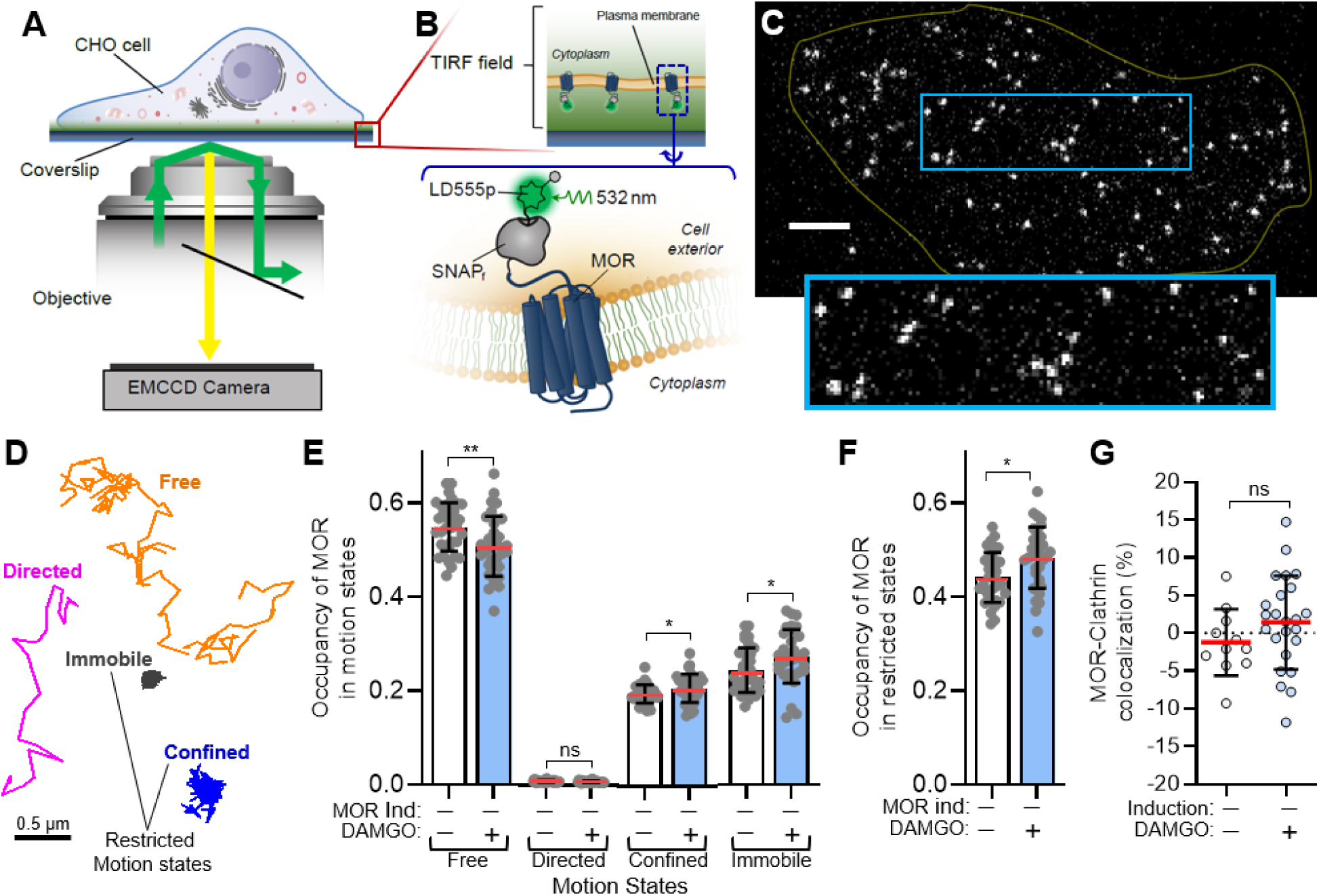
Low-density MORs show little to no engagement with clathrin-coated structures despite agonist stimulation. **A**, **B**, Schematic of single-molecule TIRF imaging of CHO cells stably expressing SNAP_f_-MOR labeled with the organic fluorophore LD555p. **C**, Representative image of a movie (frame 5, 0.075 s) of a cell excited at 532 nm in the absence of tetracycline induction showing individual unstimulated MORs in the plasma membrane. Scale bar, 5 μm; enlarged view, 5.6 μm × 20 μm; yellow line depicts the cell border. **D**, Representative single-particle trajectories showing MOR in the indicated diffusion states. **E**, Total fraction of time spent (occupancy) in the indicated motion states assigned by DC-MSS in the absence (-; white bars) (N=4, 35 cells) or presence (+; blue bars) (N=4, 34 cells) of 1 µM DAMGO. For the occupancy plots shown here and elsewhere, dots represent individual cell means and the red middle line and upper/lower lines depict the overall mean and standard deviation, respectively. **, free *p* = 0.004; not significant (ns), directed *p* = 0.2; ns, confined *p* = 0.05; *, immobile *p* = 0.02; unpaired, two-tailed t-test. **F**, Occupancy of MOR in restricted motion states, which are derived from summation of the confined and immobile states in panel E. **, *p* = 0.005, unpaired, two-tailed t-test. **G**, Fraction of individual MORs colocalized with clathrin light chain in the same cell line used in the single-particle tracking experiments in the absence (-; white symbols) (N=3, 11 cells) or presence (+; blue symbols) (N=3, 24 cells) of 1 µM DAMGO. ns, *p* = 0.2, unpaired, two-tailed t-test.

Previous studies have established that agonist-activated MORs, like many other GPCRs, localize to CCSs in a β-arrestin-dependent manner.^15,17,28–32^ This enrichment of MORs in CCSs clusters the receptors at the cell surface, and given that CCSs and their interacting receptors exhibit limited mobility at the plasma membrane prior to internalization^33^, we hypothesized that individual, agonist-treated MORs would spend more time in restricted motion states compared to untreated receptors upon assembly within CCSs. Surprisingly, treatment of uninduced cells with a saturating concentration (1 µM)^23^ of the full agonist DAMGO ([D-Ala2, N-MePhe4, Gly-ol]-enkephalin) had only a minimal effect on receptor motion states (**Figure 1E**, 0.92–1.12-fold across states), with combined restricted motion (confined + immobile) increasing by just 1.10-fold (**Figure 1F**). To directly assess receptor association with CCSs, we transiently transfected the same stable cell line with mNeonGreen-tagged clathrin light chain (CLC) and examined colocalization between SNAP_f_-labeled MOR particles and CLC in the plasma membrane (**Methods**). We observed no statistically significant DAMGO-induced colocalization under non-induction conditions, where receptor surface densities are low (**Figures 1G, S1B, and S1C**). This lack of colocalization, with both conditions yielding values near zero (−1.2% untreated, 1.4% DAMGO-treated), corroborates the minimal agonist-induced changes observed in receptor motion states. Notably, when MOR density is very low, the endogenous effectors that mediate trafficking to CCSs—β-arrestins, GRKs, adaptin, and clathrin—are expected to be in stoichiometric excess relative to receptor, yet agonist stimulation failed to measurably alter receptor motion or CCS association under these conditions.

### Sparsely labeled MORs show robust interactions with CCSs at high surface densities

Most prior studies observing agonist-induced MOR clustering within CCSs used much higher receptor densities than those employed in our experiments. To contextualize our work with these prior studies, we used the same stable MOR-expressing CHO cell line but induced receptor expression with tetracycline (**Methods**). Tetracycline induction of this cell line under similar conditions increases SNAP_f_-MOR expression by ∼500–1000-fold, reaching ∼140 ± 7 molecules/µm²,^23^ a density well above that compatible with resolving individual molecules by TIRF microscopy.

Using our inducible CHO cell line, we first sought to confirm that SNAP_f_-MORs are functional. Using a cyclic adenosine monophosphate (cAMP) inhibition assay, we demonstrated that SNAP_f_-MORs are activated in response to DAMGO treatment in both induced cells at high receptor density and, importantly, in uninduced cells at the low density used for our SPT experiments above (**Methods; Figures S2A and S2B**). These results demonstrate that the receptors are competent for G protein signaling under both uninduced and induced conditions. As expected for a downstream G protein–mediated readout, DAMGO exhibited lower apparent potency and efficacy under low-density conditions.

Having established that SNAP_f_-MORs are competent for G protein signaling, we next sought to determine whether agonist-induced trafficking to CCSs is supported at high receptor density. We transiently transfected these cells with mNeonGreen-CLC, induced MOR expression with tetracycline to achieve higher receptor density and used TIRF microscopy to assess ensemble-level clustering of SNAP_f_-MOR and colocalization with CCSs in the plasma membrane (**Methods**). Upon DAMGO treatment, we observed robust MOR clustering, with a significant fraction of these clusters colocalizing with CCSs (**Figures S2C-S2E**), consistent with prior studies of MOR^28^. Similarly, using the same cells and induction conditions, we observed strong DAMGO-stimulated MOR internalization in an enzyme-linked immunosorbent assay (ELISA) (**Methods**; **Figure S2F**), also in line with previous reports of MOR internalization following full agonist stimulation^34,35^. Thus, at high receptor density, agonist-induced clustering, CCS engagement, and endocytosis are all readily detected. In contrast, at low receptor density, MORs retain G protein signaling capacity but show minimal agonist-induced changes in motion states and no significant CCS association, highlighting a divergence between G protein–mediated signaling and GRK/β-arrestin-dependent trafficking pathways.

To relate these ensemble-level findings to our SPT experiments, we sparsely labeled tetracycline-induced SNAP_f_-MOR cells with LD555p (**Methods**; **Figure 2A**), which resulted in labeled receptor densities comparable to those achieved with more efficiently labeled receptors under non-induced low receptor expression conditions (**Figures 2B and S2G**). This allows SPT of individual labeled MORs embedded within a large excess of unlabeled receptors. Under these induction conditions, efficient labeling of SNAP_f_-MOR yielded images in which individual particles could not be resolved (**Figure 2C**), illustrating the high level of receptor expression following tetracycline induction.

**Figure 2.**
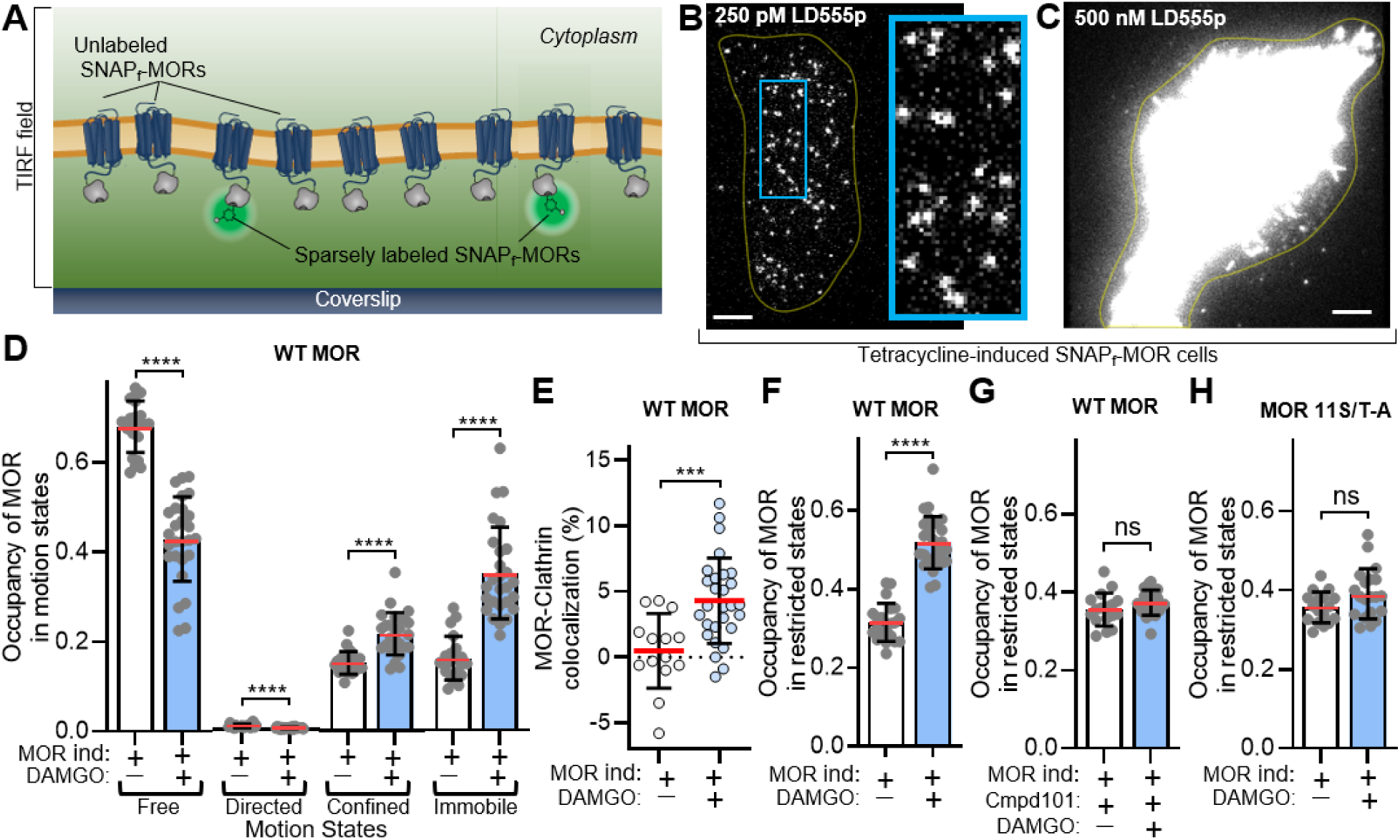
High receptor density enables robust agonist-induced MOR trafficking to clathrin-coated structures. **A**, Schematic of the highly expressed SNAP_f_-MORs in the plasma membrane of tetracycline-induced CHO cells with sparsely fluorophore-labeled receptor. **B**, **C**, Representative image of tetracycline-induced SNAP_f_-MOR stable cells labeled with (B) 250 pM or (C) 500 nM LD555p. Note that pM concentrations of the dye are needed to resolve single particles under these conditions where receptor is highly expressed. Scale bar, 5 µm. enlarged view, 5.2 μm × 18 μm; yellow line depicts the cell border. **D**, Total fraction of time spent (occupancy) in the indicated motion states assigned by DC-MSS in the absence (-; white bars) (N=3, 21 cells) or presence (+; blue bars) (N=3, 27 cells) of 1 µM DAMGO. ****: free *p* = 4 × 10^-14^; directed *p* = 5 × 10^-7^; confined *p* = 9 × 10^-7^; immobile *p* = 5 × 10^-10^; unpaired, two-tailed t-test. **E**, Fraction of individual MORs colocalized with clathrin light chain in the absence (-; white symbols) (N=3, 14 cells) or presence (+; blue symbols) (N=3, 28 cells) of 1 µM DAMGO. ***, *p* = 0.0006; unpaired, two-tailed t-test. **F**, Occupancy of MOR in restricted motion states derived from summation of the confined and immobile states in panel D. ****, *p* = 2 × 10^-15^; unpaired, two-tailed t-test. **G**, Occupancy of MOR in restricted motion states from cells treated with compound 101 (Cmpd 101) in the absence (-; white bars) (N=2, 17 cells) or presence (+; blue bars) (N=3, 27 cells) of 1 µM DAMGO. ns, p = 0.2, unpaired, two-tailed t-test. **H**, Occupancy of MOR in restricted motion states for MOR 11S/T-A in the absence (-; white bars) (N=2, 17 cells) or presence (+; blue bars) (N=2, 20 cells) of 1 µM DAMGO. ns, *p* = 0.07, unpaired, two-tailed t-test.

In contrast, DAMGO treatment of sparsely labeled SNAP_f_-MORs at high expression levels significantly reduced the time spent in the mobile spatial diffusion states and dramatically increased the time spent in the restricted motion states (**Figure 2D**). The magnitude of these changes was substantial: free state occupancy dropped to 0.63-fold of baseline, while immobile state occupancy more than doubled (2.19-fold). These results stand in stark contrast to the same experiment performed under low receptor expression conditions, where DAMGO treatment produced only very small changes in motion state occupancy (fold changes of 0.92–1.12; **Figure 1E**). Under the same high-expression, sparsely labeled conditions, DAMGO treatment significantly increased colocalization of individual MORs with mNeonGreen-CLC by nearly 9-fold (**Figures 2E, S2H, and S2I**), whereas no significant effect was observed at low receptor densities (**Figure 1G)**. These single-molecule results, combined with our bulk imaging experiments demonstrate that high-density MORs can engage CCSs despite having a vastly lower effector-to-receptor stoichiometric ratio compared to low-density conditions, where trafficking is minimal. These findings confirm that effector density is not the limiting factor in our low-density MOR experiments, since high-density MORs achieve successful engagement with fewer available effectors per receptor. Together, these data show that high receptor density is associated with robust agonist-induced CCS engagement, whereas low-density receptors exhibit minimal trafficking responses under otherwise identical conditions.

Like most other rhodopsin-like GPCRs, β-arrestin recruitment to GRK-phosphorylated MOR facilitates its interaction with CCSs. Notably, mutation of all 11 serine and threonine residues in MOR’s C-tail to alanine (MOR11S/T-A) dramatically blunts β-arrestin recruitment to the receptor and blocks its internalization^17^. To establish that the increased restricted motion states observed at high receptor density in our single-molecule experiments requires receptor phosphorylation by GRK2/3, we pretreated tetracycline-induced, sparsely labeled MOR expressing cells with the GRK2/3 inhibitor compound 101 (Cmpd101)^36^ prior to DAMGO stimulation. Unlike untreated cells where DAMGO altered mobility (**Figure 2F**), Cmpd101 treated cells showed little to no increase in restricted motion of DAMGO-activated receptors (**Figure 2G**). Additionally, unlike wild-type MOR in the same experiment, DAMGO treatment of MOR 11S/T-A—stably expressed in the same cellular background as wild-type MOR—did not increase the restricted motion states (**Figure 2H**). Together, these perturbations demonstrate that agonist-induced restricted motion at high MOR density depends on endogenous GRK2/3 activity and receptor phosphorylation.

### Nearby class A GPCRs promote MOR trafficking to CCSs

We next turned to understanding why a higher receptor density with a lower relative GRK2/3 and β-arrestin stoichiometric ratio can efficiently support the interaction of sparsely labeled MORs with CCSs. Despite unlabeled receptors outnumbering labeled MORs by ∼1000-fold, we observed robust agonist-induced CCS engagement of the labeled receptors. Rather than the abundant unlabeled MORs outcompeting the few labeled receptors for endogenous β-arrestin and GRK pools, the presence of additional activated receptors was associated with enhanced engagement of the sparsely labeled receptors with these same endogenous effectors.

To test whether the presence of additional activated GPCRs could enable CCS engagement at low MOR density, we returned to using uninduced cells. We transiently co-expressed MORs with the beta 2 adrenergic receptor (β2AR), dopamine 2 receptor (D2R), or vasopressin 2 receptor (V2R) and assessed the effect on MOR mobility (**Methods**). Like MOR, β2AR and D2R are broadly characterized as class A β-arrestin-interacting receptors from which β-arrestin dissociates after internalization^37,38^. In contrast, V2R is a prototypical class B β-arrestin-interacting receptor that forms stable complexes with β-arrestin that persist after internalization^37,39^ (Here, the terms class A and class B refer specifically to β-arrestin binding affinity and internalization behavior, and not to the classical class A–F GPCR classification based on sequence homology^40,41^). The transiently expressed β2AR, D2R and V2R contain N-terminal HaloTags^42^ for orthogonal labeling with JF646^43^ (**Figure 3A**). Halo-tagged receptor expression was dramatically higher than MOR levels in uninduced cells and therefore not suitable for SPT (**Figures 3B and S3A**). Importantly, GPCR co-expression did not affect MOR expression levels (**Figures S3B-S3D).**

**Figure 3.**
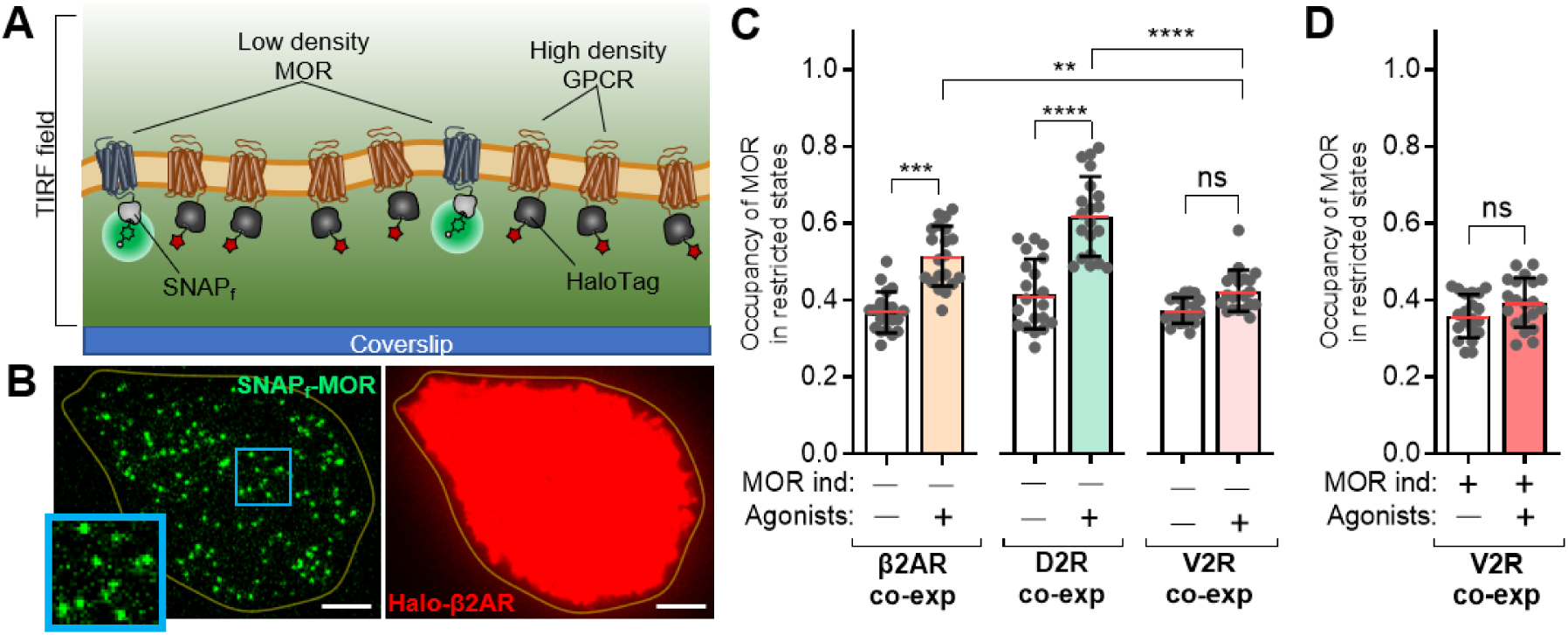
Co-expressed class A GPCRs enhance MOR trafficking to CCSs, whereas V2R suppress it. **A**, Schematic of low density and efficiently labeled SNAP_f_-MORs coexpressed with Halo-tagged GPCRs at higher receptor densities in the plasma membrane of CHO cells. **B**, (left) Representative image from a movie (frame 5, 0.075 s) of an uninduced cell excited by 532 nm showing individual MORs in the plasma membrane and (right) an image of the same cell taken prior to the movie excited by 640 nm showing β2AR coexpression at the cell surface. Scale bar, 5 μm; enlarged view, 5.3 μm × 5.1 μm; yellow line depicts the cell border. **C**, Occupancy of MOR in restricted motion states from uninduced cells with the indicated co-expressed neighboring receptors in the absence (-) or presence (+) of 1 µM DAMGO and full agonists for the co-expressed receptors (10 µM isoproterenol for β2AR; 10 µM quinpirole for D2R; 250 pM arginine vasopressin for V2R). For all experiments, N=2, 20 cells. ****, *p* = 4.8 x 10^-10^; ***, *p* = 4.5 x 10^-8^; **, *p* = 0.0015; ns, *p* = 0.20. Multiple comparisons between the three experiments in the absence of agonists (white bars) were all not significant (not indicated on plot) with *p* ≥ 0.41. Two-way ANOVA with Tukey multiple comparisons test. **D**, Occupancy of high density, sparsely labeled MOR in restricted motion states from tetracycline-induced cells co-expressed with V2R in the absence (-) (N=2, 20 cells) or presence (+) (N=2, 20 cells) of 1 µM DAMGO and 250 pM arginine vasopressin. ns, p = 0.1, unpaired, two-tailed t-test.

In the corresponding SPT experiments, DAMGO stimulation significantly increased MOR restricted motion states when β2AR and D2R were co-expressed and co-stimulated with their full agonists, isoproterenol and quinpirole, respectively (**Figure 3C**). The magnitude of these agonist-induced changes—1.40-fold for β2AR and 1.49-fold for D2R co-expression—approached that observed when MOR alone was highly expressed (1.64-fold; **Figure 2F**) and far exceeded the smaller fold changes observed for MOR at low density without co-expression (1.1-fold; **Figure 1F**). These results suggest that high-density class A receptors, such as β2AR and D2R increase the probability of productive engagement between endogenous GRK2/3 and β-arrestin and sparsely expressed MORs, promoting MOR engagement with CCSs.

Intriguingly, unlike the experiments with co-expressed β2AR and D2R, co-expression of the high-affinity β-arrestin-interacting V2R with its full agonist produced no significant change in MOR restricted motion (**Figure 3C**), with the small 1.14-fold increase closely matching the minimal 1.10-fold change for MOR alone at low density (**Figure 1F**). We note that the relative expression of V2R was similar to β2AR and D2R (**Figures S3B-D**), suggesting that this observation is not related to differences in surface expression between these receptors. Tetracycline-induced cells with highly expressed, sparsely labeled MOR showed large DAMGO-stimulated increases in MOR restricted motion states in the absence of GPCR coexpression (**Figures 2D and 2F**). Remarkably, this effect was abolished when V2R was co-expressed and co-stimulated (**Figure 3D**). These data indicate that highly expressed V2R does not support MOR restricted motion in the same manner as class A β-arrestin–interacting receptors and instead suppresses agonist-induced MOR restricted motion at both low and high MOR expression levels.

### Increased effector density near the plasma membrane promotes MOR engagement with CCSs

If abundant class A GPCRs create higher effective availability of β-arrestins and GRK2/3 near the membrane, we hypothesized that increasing the density of these effectors in the absence of additional transfected receptors might mimic this effect in low-density MOR cells. To increase the density of these effectors, we transiently expressed β-arrestin2 or GRK2 fused to HaloTags (**Figure 4A**) in uninduced cells where SNAP_f_-MOR densities are low. Like our receptor co-expression experiments, these cells were labeled with HaloTag-reactive, cell-permeant JF646 to select those containing the expressed effectors and with concentrations of SNAP_f_-reactive LD555p for efficient labeling of MORs (**Methods**). The receptor densities remained at single-particle resolution and similar to untransfected cells (**Figures 4B and S4A-S4C**). Brief excitation of the same cells at the JF646 wavelength showed robust fluorescence in the TIRF imaging field (**Figures 4B and S4C**), consistent with increased effector abundance within the evanescent field and therefore in close proximity to the plasma membrane. Like our class A GPCR co-expression results, DAMGO treatment of these cells increased the fraction of time low-density MORs spent in the restricted motion states by 1.4- and 1.5-fold (**Figures 4C and 4D**), demonstrating that higher effector abundance enhances the agonist-induced occupancy of MOR restricted states, consistent with CCS engagement.

**Figure 4.**
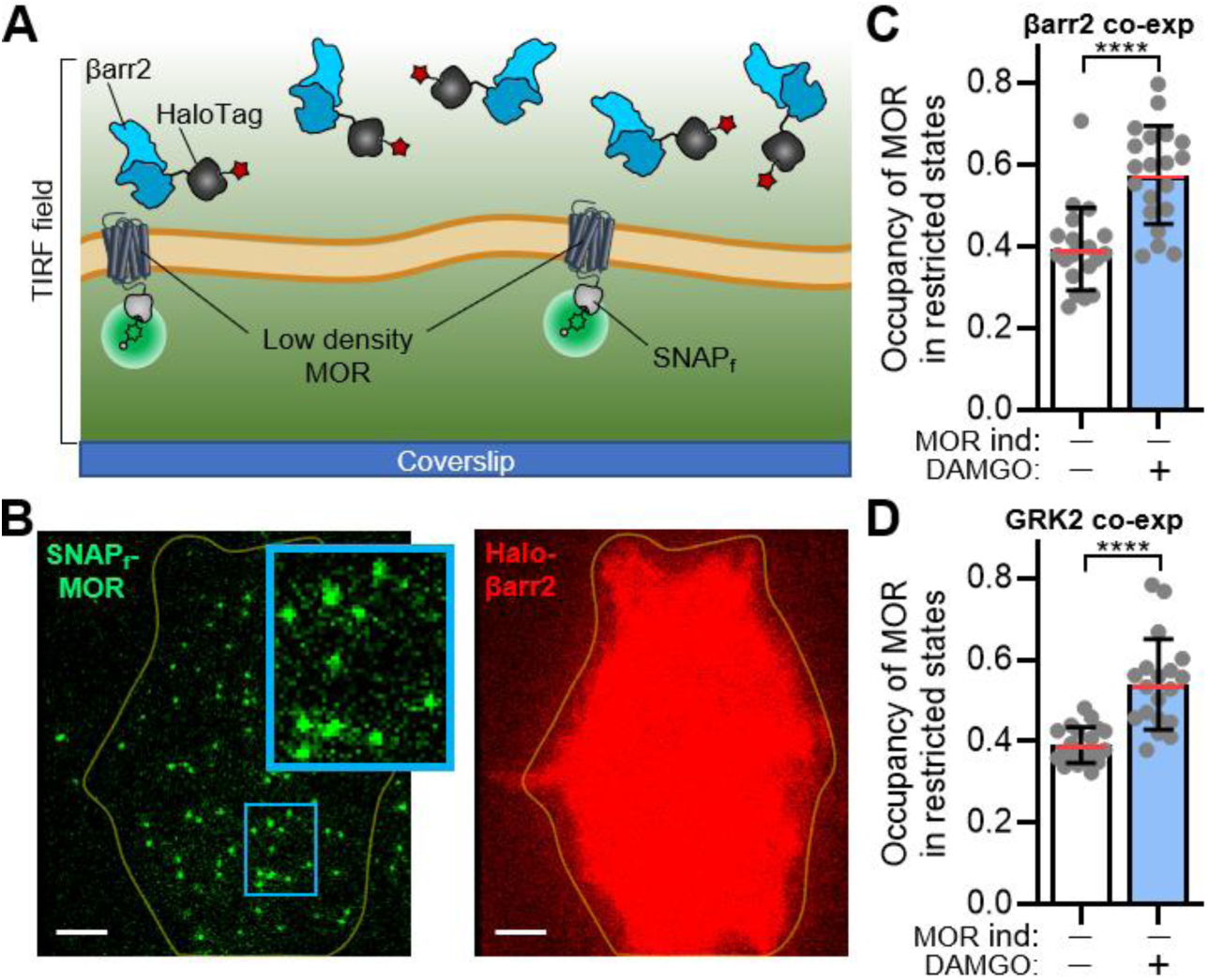
Increasing β-arrestin2 or GRK2 expression enhances MOR engagement with CCSs at low receptor densities. **A**, Schematic of the low density, efficiency labeled SNAP_f_-MORs in the plasma membrane of uninduced CHO cells with co-expressed Halo-β-arrestin2 (βarr2) (shown in schematic) or Halo-GRK2 (not shown). **B**, (left) Representative image from a movie (frame 5, 0.075 s) of an uninduced cell excited by 532 nm showing individual MORs in the plasma membrane and (right) an image of the same cell taken prior to the movie excited by 640 nm showing intracellular βarr2. Scale bar, 5 μm; enlarged view, 3.3 μm × 5.1 μm; yellow line depicts the cell border. **C**, Occupancy of MOR in restricted motion states from cells with coexpressed βarr2 in the absence (-; white bars) (N=2, 20 cells) or presence (+; blue bars) (N=2, 20 cells) of 1 µM DAMGO. ****, *p* = 8 × 10^-6^, unpaired, two-tailed t-test. **D**, Occupancy of MOR in restricted motion states from cells with coexpressed GRK2 in the absence (-; white bars) (N=2, 20 cells) or presence (+; blue bars) (N=2, 19 cells) of 1 µM DAMGO. ****, *p* = 2 × 10^-6^, unpaired, two-tailed t-test.

## DISCUSSION

Using live-cell single-molecule imaging, we demonstrate that high-density MORs facilitate the trafficking of sparsely labeled MORs to CCSs, a process that cannot be efficiently engaged when MORs are expressed at low density alone. One mechanism that might explain this increased effector efficiency is density-dependent homodimerization between sparsely labeled and unlabeled MOR protomers. However, we recently showed using the same cells that SNAP_f_-MOR is predominantly monomeric under both the low and high density conditions used in this study,^23^ ruling out density-dependent dimerization as a contributor to this mechanism. We propose instead that the abundant unlabeled MORs serve as the major driver of endogenous β-arrestin and/or GRK2/3 recruitment, forming an “affinity matrix” for cytosolic effectors—a receptor-driven, density-dependent platform that supports reversible interactions without net sequestration—thereby increasing the probability of productive effector interactions with sparsely labeled receptors. (**Figures 5A-5D**).

**Figure 5.**
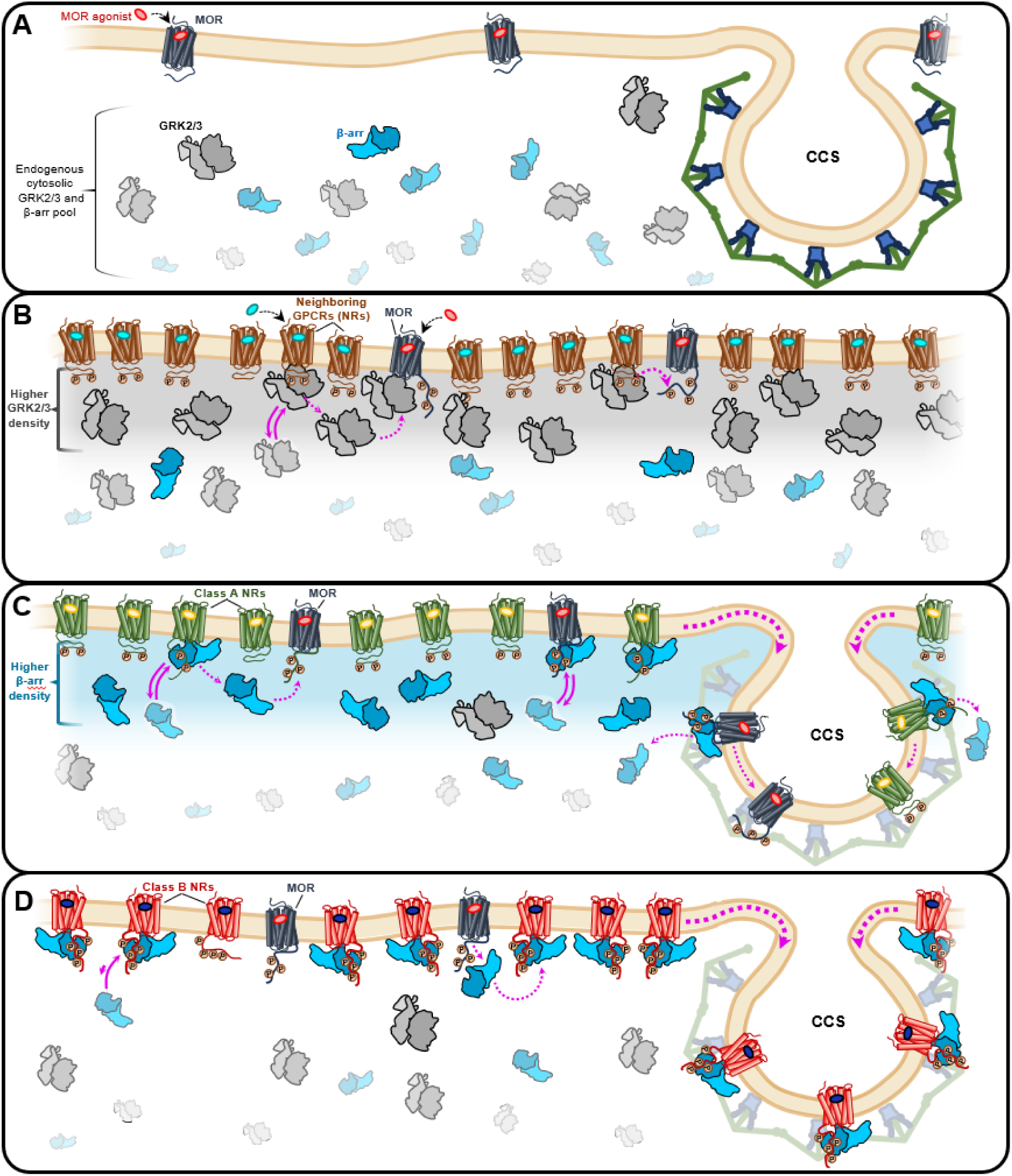
Model for density-dependent regulation of μ-opioid receptor trafficking by class A and class B GPCRs. **A,** At low receptor density, MOR activates G proteins but fails to efficiently engage GRK2/3 and β-arrestin, resulting in minimal trafficking to clathrin-coated structures (CCSs). **B,C,** As the density of class A GPCRs increases, despite worsening effector-to-receptor stoichiometry, activated receptors form an affinity matrix that enables reversible GRK2/3 and β-arrestin interactions to be productively used by neighboring receptors, permitting MOR phosphorylation, β-arrestin engagement, and trafficking to CCSs. **D,** In contrast, class B GPCRs such as V2R bind β-arrestin with high affinity and act as sequestration sinks, monopolizing β-arrestin availability and blocking GRK2/3–β-arrestin–dependent trafficking of nearby MORs. This model summarizes how receptor density, rather than intrinsic receptor competence, tunes signaling pathway engagement and trafficking outcomes.

The findings of two prior single-molecule studies of MOR seem consistent with our proposal of density-dependent CCS engagement. Metz et al. observed large increases in receptor immobility and clathrin colocalization after DAMGO treatment using high-expression AtT20 cells with sparse Qdot labeling,^44^ while Gentzsch et al., working at moderate expression levels (∼0.8 particles/μm²), found that only ∼23% of receptor immobilization events overlapped with clathrin-coated pits^45^. Here, by systematically varying MOR density within the same experimental system, we directly isolate receptor density as a key variable governing MOR engagement with CCSs.

Our experiments co-expressing other class A receptors at high densities with low-density MOR further support this model: like the unlabeled MORs, these neighboring receptors drive effector recruitment at high density. Because activated GPCRs engage GRKs through transient interactions,^46^ neighboring receptors have the potential to support efficient GRK2/3-dependent phosphorylation and downstream β-arrestin engagement for nearby MORs. Critically, β2AR and D2R, like MOR, are class A GPCRs with lower β-arrestin affinity,^37,38^ allowing them to recruit and release β-arrestin rather than sequester it, establishing an affinity matrix that promotes MOR trafficking to CCSs (**Figures 5B and 5C**).

Overexpression of β-arrestin2 or GRK2 resulted in robust increases in restricted-state motion of low-density MORs, further supporting the idea that increasing effector availability near the plasma membrane can facilitate MOR engagement with CCSs. Agonist-induced restricted motion requires both GRK2/3 activity (Cmpd101 sensitivity) and MOR C-tail phosphorylation sites (MOR11S/T-A). Thus, increased GRK2 availability most likely enhances the efficiency of phosphorylating sparsely expressed MORs^47,48^, promoting downstream β-arrestin–dependent engagement with CCSs. Importantly, overexpression of β-arrestin2 alone was also sufficient to rescue agonist-induced interactions of low-density MORs with CCSs, consistent with a model in which increased β-arrestin availability enhances the likelihood that MORs engage β-arrestin and enter CCS-associated restricted states. It is also possible that a fraction of overexpressed β-arrestin preassociates directly with the plasma membrane,^49^ further facilitating receptor engagement through lateral diffusion. Regardless of the precise mechanism, the failure of endogenous β-arrestin to support trafficking at low receptor density underscores that productive receptor–β-arrestin engagement is fundamentally governed by encounter probability, which at endogenous β-arrestin levels requires sufficient receptor density.

At endogenous effector levels, we propose that neighboring receptors contribute to GRK transphosphorylation, i.e. phosphorylation of one receptor by GRKs recruited by neighboring receptor activation. Indeed, the neuropeptide FF2 receptor was shown to phosphorylate residues on MOR when co-expressed with MOR.^50^ While this transphosphorylation was attributed to GPCR heterodimerization, we propose a different mechanism in which physical and stable interactions between receptors are not required. Instead, activation of neighboring receptors in close proximity generates increased local densities of GRK2/3, allowing these kinases to be shared between like and unlike receptor types.

Unlike β2AR and D2R, coexpression of V2R, a prototypical class B GPCR, with MOR at both low and high densities blocked MOR interaction with CCSs. A likely mechanism is V2R’s high affinity for β-arrestin, which allows it to sequester β-arrestin from MOR and/or compete for β-arrestin already engaged with MOR (**Figure 5D**). Thus, whereas class A β-arrestin-interacting GPCRs function as an affinity matrix that supports reversible, non-sequestering access to GRK2/3 and β-arrestin, V2R and presumably other class B GPCRs act as sequestration sinks that monopolize β-arrestin availability. Such mechanisms are not unique to β-arrestin, as a prior study has shown that a neighboring receptor can sequester G protein and interfere with G protein activation of other GPCRs.^51^

Our model of an affinity matrix is compatible with the catalytic activation mechanism described by Eichel et al., in which β-arrestin can remain in an active conformation at the membrane after dissociating from class A receptors.^52^ Catalytic activation may contribute to the pool of β-arrestin available for receptor engagement, and the two mechanisms are not mutually exclusive. However, catalytic activation alone is unlikely to fully account for our observations. Even if β-arrestin is catalytically activated by sparse receptors, it must still productively engage phosphorylated receptors to mediate trafficking—and our data demonstrate that this engagement is inefficient at low receptor density despite a large stoichiometric excess of endogenous β-arrestin. We propose that high receptor density is required to capture and retain β-arrestin—whether newly recruited or catalytically activated—through an affinity matrix that increases the probability of productive engagement.

Our findings reveal an important distinction in how receptor density regulates different signaling pathways downstream of MOR. While G protein signaling remains functional at both low and high MOR densities, the GRK2/3-β-arrestin-mediated pathway, and consequently receptor trafficking to CCSs, exhibits density dependence. At low receptor densities, MORs can activate G proteins but fail to efficiently recruit GRK2/3 or β-arrestin and traffic to CCSs, effectively favoring G protein signaling at the plasma membrane. Because G proteins are lipid-anchored and already membrane-localized,^53^ their interaction with MOR is less dependent on local recruitment than that of cytosolic β-arrestin and GRK2/3. In contrast, at high receptor densities where an affinity matrix for cytosolic effectors is established, both the G protein and GRK2/3-β-arrestin pathways are able to operate concurrently. This density-dependent biasing represents a fundamentally different regulatory mechanism for receptor function, where the spatial organization and local concentration of receptors, in addition to receptor conformation or ligand properties, together determine the signaling outcome.

GPCRs have previously been shown to influence the internalization of other GPCR subtypes with these effects often attributed to GPCR heterodimerization.^54,55^ However, our observations with MOR alone, which does not homodimerize,^23^ cannot be explained by physical receptor-receptor association. Rather, whether a co-expressed GPCR facilitates or prevents MOR trafficking depends on its β-arrestin binding class: class A receptors promote effector sharing, while class B receptors such as V2R sequester β-arrestin. Consistent with this, the class B GPCR Neurokinin 1 receptor was shown to inhibit endocytosis of MOR, an effect overcome by the addition of β-arrestin.^56^ Thus, rather than heterodimerization,^57^ density-dependent effector sharing or sequestration provides an alternative mechanism by which GPCRs can regulate each other’s trafficking.

Because efficient affinity matrix formation requires sufficiently high receptor densities, the local density of MOR and other class A GPCRs will fundamentally determine trafficking responses in physiologically relevant contexts. Indeed, MOR has been shown to internalize in dendrites but not in axon terminals in response to morphine,^58,59^ raising the possibility that differences in local receptor density across neuronal compartments contribute to these distinct trafficking outcomes. In highly compartmentalized cells like neurons, receptor and effector densities likely vary across diverse structures such as dendritic spines, axon terminals, and somatic regions. The many scaffolding proteins that can interact with GPCRs and effectors, sometimes in a microdomain-specific manner, will also enable differences in function. Importantly, GPCRs are highly co-expressed across the brain, and many GPCR subtypes can be present in the same neuron^60,61^. Here, we demonstrate a mechanism by which one activated GPCR can potentiate or suppress the trafficking of another GPCR type. Taken together, the local density of MOR, other activated GPCRs, and shared cytosolic effectors will ultimately determine the signaling outcome. These findings may also be relevant to polypharmacological approaches in which multiple GPCRs are targeted simultaneously,^1,11,62^ as the density-dependent crosstalk we describe provides a mechanism by which activation of one receptor could influence the trafficking and signaling of another.

### Limitations of the study

We used CHO cells as a model system, which lack the complex membrane organization and scaffolding proteins present in neurons where MORs naturally function. While this simplified system enabled precise control of receptor density, the mechanisms we describe may be modified by cell-type-specific factors and the diverse scaffolding environments found in native tissues. Our co-expression experiments relied on transient transfection to achieve high GPCR densities, which may not fully recapitulate physiological expression patterns or stoichiometries observed *in vivo*. We also primarily used DAMGO as a full agonist. Different agonists with varying efficacy profiles may produce distinct density-dependent effects on trafficking. Additionally, while we infer changes in the effective availability of endogenous GRK2/3 and β-arrestin from receptor behavior, we did not directly visualize the spatial distributions or dynamics of these endogenous effectors under different receptor densities.

## RESOURCE AVAILABILITY

### Lead contact

Requests for further information or resources and reagents should be directed to and will be filled by the lead contact, Jonathan A. Javitch.

### Materials availability

Plasmids and cell lines generated in this study are available from the lead contact with a completed material transfer agreement.

## DATA AND CODE AVAILABILITY

- Source data generated and analyzed that support the findings of this study will be shared by the lead contact upon request.
- This paper does not report original code. All software used to collect and analyze data for this work was either published previously or is commercially available.
- Any additional information required to reanalyze the data reported in this paper is available from the lead contact upon request.

## ACKNOWLEDGEMENTS

This work was supported by NIH grants R01 MH054137 (J.A.J.), R35 GM145284 (N.L.), the Hope for Depression Research Foundation (J.A.J.), the St. Jude Children’s Research Hospital GPCR Collaborative (JAJ), Miriam’s Magical Memorial Mission (J.A.J.), the Brain and Behavior Research Foundation NARSAD Young Investigator Award (W.B.A.) and the Charles Revson Foundation/Burroughs Wellcome Fund PDEP Fellowship (C.M.W). We thank Marina Dawoud and Lisa Lin for technical assistance. We thank Robert J. Lane for helpful discussions. We thank Scott C. Blanchard for providing BG-LD555p.

## AUTHOR CONTRIBUTIONS

M.D.H and W.B.A. prepared an initial draft outlining the work, and W.B.A and J.A.J. wrote and edited the final submitted manuscript, with contributions from all the authors. M.D.H, P.G., W.B.A., and J.A.J designed the single-molecule experiments and interpreted the results with critical input from M.C and N.A.L. All of the live-cell single-molecule imaging experiments of MOR diffusion were performed by M.D.H and the resulting data were analyzed by M.D.H, P.G., C.M.W. and W.B.A., with input related to tracking and DCMSS analysis optimization from S.J.M. and P.G. and global input by W.B.A. and J.A.J. Select elements of the single-molecule data analysis were confirmed by C.M.W. and P.G. with input from J.A.J and W.B.A. The single-molecule and bulk receptor-clathrin colocalization experiments were performed and analyzed by A.B., with input by W.B.A., M.D.H, J.A.J. and N.A.L. The cell-based ELISA MOR internalization assay was performed by A. G. and W.B.A. and analyzed by W.B.A. The cAMP assays were performed and analyzed by G.V.D. with critical input from J.A.J and W.B.A. The overall project was supervised by W.B.A and J.A.J.

## DECLARATION OF INTERESTS

### Declaration of generative AI and AI-assisted technologies in the writing process

During the preparation of this work, the authors used Claude AI to improve the readability and language of the manuscript. The authors reviewed and edited all AI-generated content and take full responsibility for the content of the published article.

## SUPPLEMENTAL INFORMATION FIGURE TITLES AND LEGENDS

**Figure S1.**
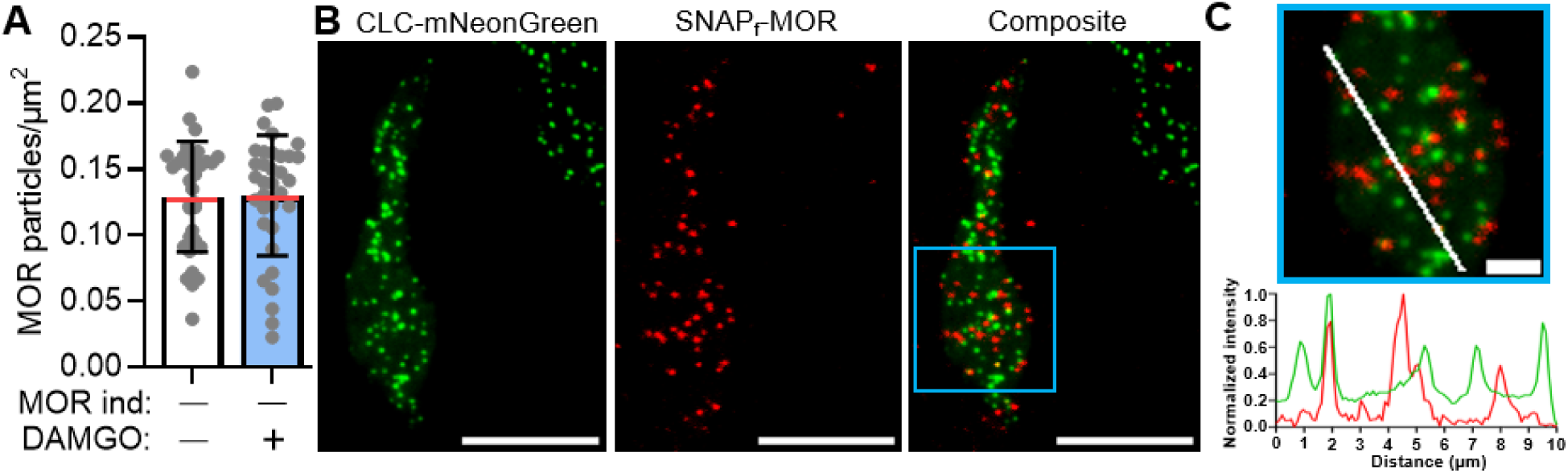
SNAP_f_-MOR surface density in uninduced cells is compatible with single-particle tracking and clathrin colocalization analysis. **A**, Labeled surface density of SNAP_f_-MORs in uninduced, efficiently labeled cells in the absence (-; white bar) (N=4, 35 cells) or presence (+; blue bar) (N=4, 34 cells) of 1 µM DAMGO (determined in frame 5 of movies). For the receptor density plots shown here and elsewhere, dots represent individual cell means of the number of MOR particles per cell area and the red middle line and upper/lower lines depict the overall mean and standard deviation, respectively. **B**, Representative image of a cell treated with 1 µM DAMGO showing clathrin light chain (CLC)-mNeonGreen (left), SNAP_f_-MOR (middle), and the composite of the two (right) showing lack of colocalization. Blue box in the composite image denotes the region shown at higher magnification in (C). Scale bar, 10 µm. The analysis of multiple cell images shown in Figure 1G. **C**, (top) Enlarged view of the boxed region in (B) with a white line indicating the path of the fluorescence intensity line profile. Scale bar, 2 µm. (bottom) Fluorescence intensity line profile along the white line in the enlarged image showing the normalized intensity of CLC-mNeonGreen (green) and SNAPf-MOR (red) as a function of distance (µm), illustrating the lack of spatial overlap between individual MOR particles and CLC puncta at low receptor density.

**Figure S2.**
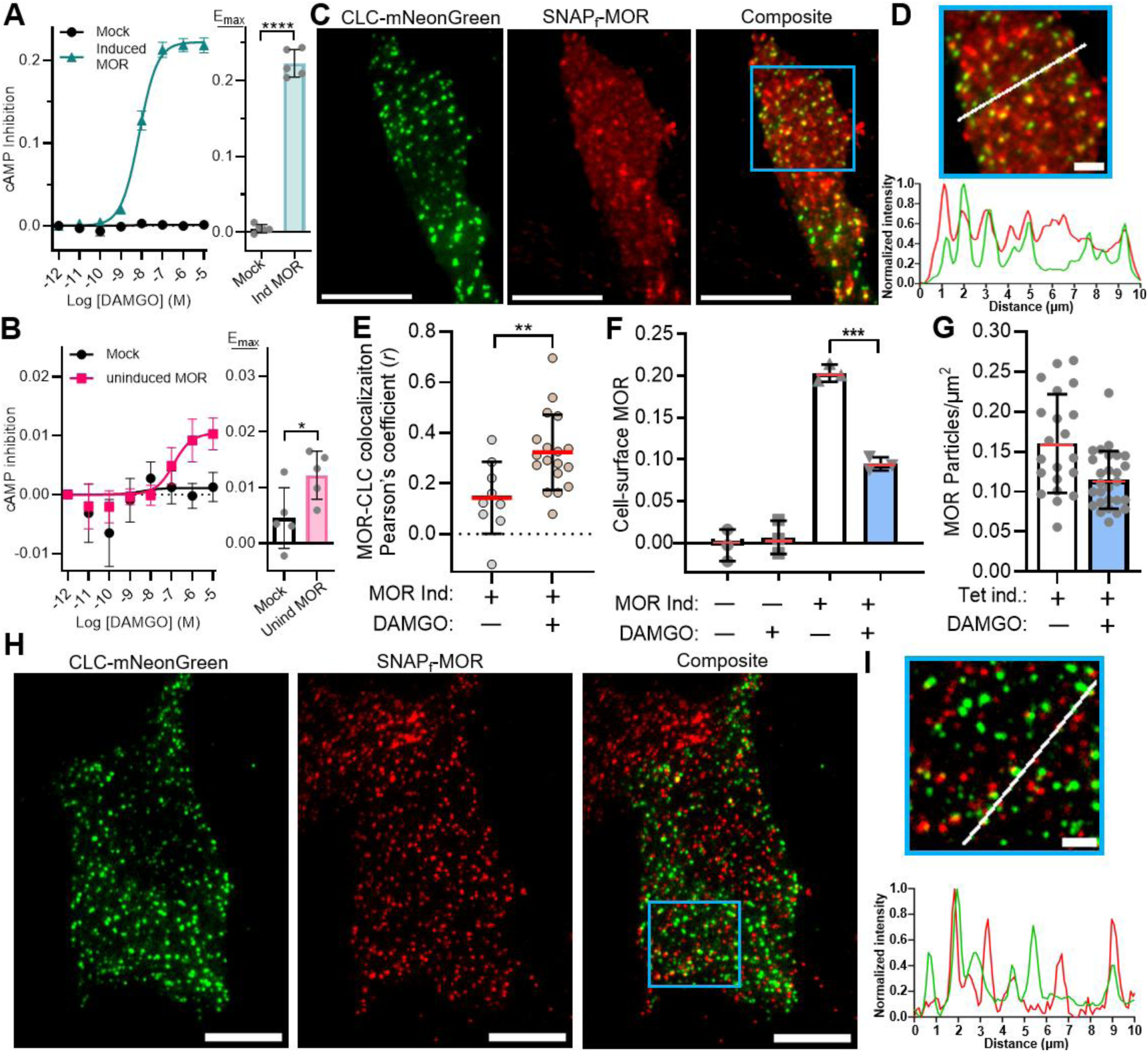
SNAP_f_-MORs are functional for G protein signaling and exhibit agonist-induced clathrin colocalization and internalization at high density. **A**, **B**, (left) cAMP-inhibition measurements showing DAMGO-concentration-response curves (CRCs) and (right) corresponding E_max_ distributions for CHO cells not expressing MOR (mock) and the CHO cells stably expressing SNAP_f_-MOR used in our imaging experiments in the (A) presence (induced MOR) and (B) absence (uninduced MOR) of tetracycline induction. For the CRC plots, symbols represent the mean drug-induced BRET response and error bars represent ± SEM for N=5 independent experiments performed in triplicate. For the E_max_ plots, symbols represent the mean E_max_ value for each experiment and error bars represent ± SD. ****, *p* = 6 × 10⁻⁹; *, *p* = 0.03 unpaired; two-tailed t-test. **C**, Representative image of a cell treated with 1 µM DAMGO showing clathrin light chain (CLC)-mNeonGreen (left), SNAP_f_-MOR (middle), and the composite of the two (right) showing receptor colocalization with CLC. Blue box in the composite image denotes the region shown at higher magnification in (D). Scale bar, 10 µm. **D**, (top) Enlarged view of the boxed region in (C) with a white line indicating the path of the fluorescence intensity line profile. Scale bar, 2 µm. (bottom) Fluorescence intensity line profile along the white line in the enlarged image showing the normalized intensity of CLC-mNeonGreen (green) and SNAP_f_-MOR (red) as a function of distance (µm), illustrating spatial overlap between MOR clusters and CLC puncta following DAMGO treatment at high receptor density. **E**, Quantification of SNAP_f_-MOR colocalization with CLC-mNeonGreen as Pearson’s coefficient (*r*) for many cells in the absence (−; gray symbols) (N=3, 9 cells) and presence (+; brown symbols) (N=3, 18 cells) of 1 µM DAMGO. Here and elsewhere, dots represent individual cell means and the middle and upper/lower lines depict the overall mean and standard deviation, respectively. **, *p* = 0.0061, unpaired, two-tailed t-test. **F**, ELISA-based receptor internalization assay using the same stable CHO cell line used in the single-molecule experiments expressing SNAP_f_-MOR in the absence (−) or presence (+) of tetracycline induction as well as in the absence (−; white bars) or presence (+; blue bars) of 1 µM DAMGO treatment. The MOR surface signal is representative of the background-corrected absorbance (OD₄₉₀). In the absence of tetracycline induction, the signal is indistinguishable from background due to low MOR expression levels that can only be detected with single-molecule methods. ***, *p* = 0.0001, unpaired, two-tailed t-test. **G**, Surface density of SNAP_f_-MORs in tetracycline-induced, sparsely labeled cells in the absence (−; white bar) or (+; blue bar) presence of 1 µM DAMGO (determined in frame 5 of movies). **H**, Representative image of a cell after 1 µM DAMGO treatment showing clathrin light chain (CLC)-mNeonGreen (left), SNAP_f_-MOR (middle), and the composite of the two (right) showing colocalization. Blue box in the composite image denotes the region shown at higher magnification in (I). Scale bar, 10 µm. The analysis of multiple cell images shown in Figure 2E. **I**, (top) Enlarged view of the boxed region in (H) with a white line indicating the path of the fluorescence intensity line profile. Scale bar, 2 µm. (bottom) Fluorescence intensity line profile along the white line in the enlarged image showing the normalized intensity of CLC-mNeonGreen (green) and SNAP_f_-MOR (red) as a function of distance (µm), illustrating spatial overlap between sparsely labeled MOR particles and CLC puncta at high receptor density.

**Figure S3.**
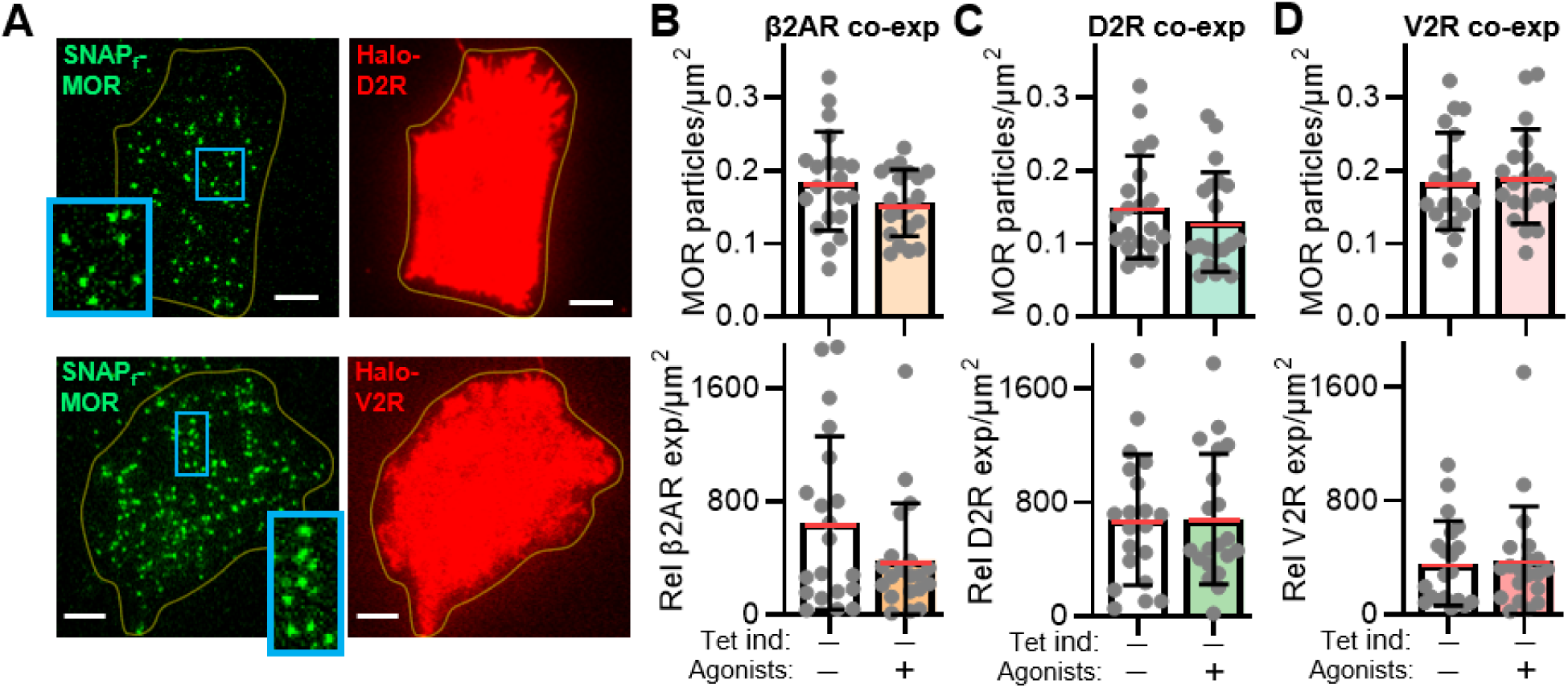
The cell-surface density of SNAP_f_-MOR in uninduced cells is unaffected by coexpression of neighboring GPCRs. **A**, (left) Representative image from a movie (frame 5, 0.075 s) of an uninduced cell excited by 532 nm showing individual MORs in the plasma membrane and (right) an image of the same cell taken prior to the movie excited by 640 nm showing (top) D2R and (bottom) V2R coexpression at the cell surface. Scale bar, 5 μm; top enlarged view, 5.0 μm × 5.8 μm; bottom enlarged view, 3.4 μm × 6.7 μm; yellow line depicts the cell border. **B**, **C**, **D**, (top) Labeled surface density of SNAP_f_-MORs in uninduced, efficiently labeled cells co-expressing (B) β2AR, (C) D2R, or (D) V2R and (bottom) the corresponding relative β2AR, D2R, and V2R expression in the absence (-; white bar) or presence (+; orange for β2AR, green for D2R, and pink for V2R) of 1 µM DAMGO (determined in frame 5 of movies). For all plots, N=2, 20 cells. For the MOR density plots, dots represent the number of MOR particles per cell area. For the relative neighboring GPCR expression plots, dots represent the total background-corrected fluorescence per cell area.

**Figure S4.**
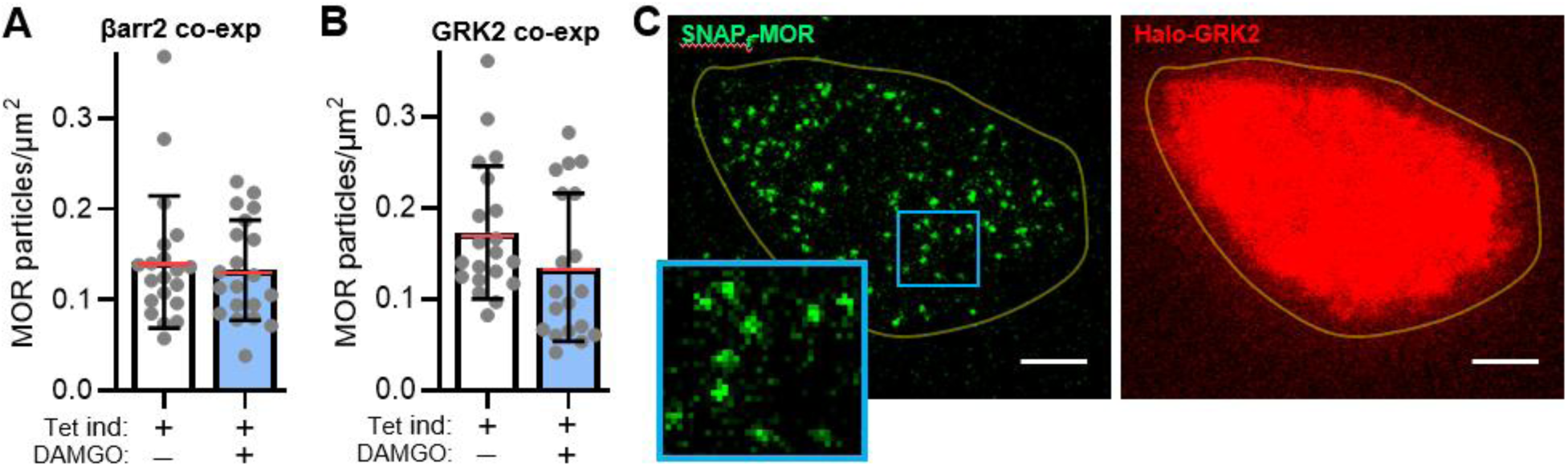
The cell-surface density of SNAP_f_-MOR in uninduced cells is unaffected by coexpression of β-arrestin2 or GRK2. **A**, Labeled surface density of SNAP_f_-MORs in uninduced, efficiently labeled cells co-expressing Halo-β-arrestin2 in the absence (-; white bar) (N=2, 20 cells) or presence (+; blue bar) (n=N, 20 cells) of 1 µM DAMGO (determined in frame 5 of movies). **B**, Labeled surface density of SNAP_f_-MORs in uninduced, efficiently labeled cells co-expressing Halo-GRK2 in the absence (-; white bar) (N=2, 20 cells) or presence (+; blue bar) (N=2, 19 cells) of 1 µM DAMGO (determined in frame 5 of movies). **C**, (left) Representative image from a movie (frame 5, 0.075 s) of an uninduced cell excited by 532 nm showing individual MORs in the plasma membrane and (right) an image of the same cell excited by 640 nm taken prior to the movie showing intracellular Halo-GRK2. Scale bar, 5 μm; enlarged view, 3.8 μm × 4.0 μm; yellow line depicts the cell border.

## SUPPLEMENTAL INFORMATION TABLE AND LEGEND

**Table S1.** Tracking parameters within u-track used for single-particle tracking.

| Input parameters | Value |
| --- | --- |
| *Spot detection parameters* |  |
| Psf-sigma | 0.7 pixels |
| Mixture-model fitting | 1 |
| Alpha value for initial detection of local maxima | 0.015 |
| Integration window | 3 |
| **testAlpha R, A, D, F** | 0.05, 0.05, 0.05, 0 |
| *General tracking parameters* |  |
| Time window | 5 frames |
| **Flag for merging and splitting** | 0 |
| Minimum track segment length | 2 frames |
| *Frame-to-frame linking parameters* |  |
| Flag for linear motion | 0 |
| Search radius lower limit | 2 pixels |
| Search radius upper limit | 3 pixels |
| Std dev multiplication factor | 3 |
| Flag for using local density | 1 |
| Number of frames used in nearest neighbor calculation | 5 |
| *Gap closing, merging and splitting parameters* |  |
| Flag for linear motion | 0 |
| Search radius lower limit | 2 pixels |
| Search radius upper limit | 3 pixels |
| Std dev multiplication factor | [3 3 3 3 3] |
| Brownian scaling | [0.25, 0.25] |
| timeReachConfB | 5 |
| Amplitude ratio lower and upper limit | [0.7, 4] |
| Min length for track segment analysis | 5 |
| Flag for using local density | 0 |
| Number of frames used in nearest neighbor calculation | 5 |
| linStdMult | [1 1 1 1 1] |
| linScaling | [0.25 0.25] |
| timeReachConfL | 5 |
| Max angle between the directions of motion | 45 |
| Gap penalty | 1.5 |
| gapExcludeMS | 1 |
| strategyBD | -1 |

## METHODS

### EXPERIMENTAL MODEL AND SUBJECT DETAILS

#### CHO cell culture

All CHO lines were maintained in Ham’s F12 medium (Corning), 10% FBS (Corning), 2 mM l-glutamine (Corning) and 1% penicillin–streptomycin at 37 °C in 5% CO2. For LEx-Flp-In T-REx (FITR) CHO cells, the medium contained 15 μg ml^−1^ blasticidin (InvivoGen) and 50 μg ml^−1^ zeocin (Invitrogen). For the SNAP_f_-MOR stable LEx-FITR CHO line, the medium contained 500 μg ml^−1^ hygromycin (Thermo Fisher Scientific) and 15 μg ml^−1^ blasticidin. All CHO lines were grown to ∼70% confluency in six-well tissue culture dishes before transfection, tetracycline (MilliporeSigma) induction and labeling for microscopy, ELISA, and other experiments. All cell lines used in this study tested negative for mycoplasma contamination. CHO-K1 cells, the parent line of our CHO cells used in this study, do not natively express MOR based on an online database of CHO RNA-seq data.^63^

## METHOD DETAILS

### Reagents and organic fluorophores

LD555p-benzylguanine (BG) was prepared from LD555p-NHS (Lumidyne Technologies) as previously described.^64^ Halo-JF646 was obtained from Luke D. Lavis (Jamelia Research Campus, Howard Hughes Medical Institute). DY549-BG was purchased from New England Biolabs.

### Plasmid construction

The pcDNA5/FRT/TO-IRES vector encoding amino-terminally SNAP_f_-tagged MOR (SNAP_f_-MOR) was constructed as previously described^23^ by replacing the dopamine D2 receptor coding region of the pcDNA5/FRT/TO-IRES SNAPfast-D2 vector with the human MOR coding region using standard subcloning methods. The phosphorylation-deficient mutant SNAPf-MOR11S/T-A was cloned into the same pcDNA5/FRT/TO-IRES vector. In this vector, the Kozak sequence was mutated to thymines at the −3 and +4 positions to produce a weak Kozak sequence, and the CMV enhancer sequence was partly deleted by digestion with NruI and SnaBI and subsequent religation to generate a crippled CMV promoter, both of which contribute to low basal expression levels. All constructs were sequence-verified (Macrogen). For co-expression experiments, Halo-tagged constructs were cloned into pcDNA3.1 (β2AR, V2R, GRK2), pcDNA3.0 (GRK2-Halo), or pIRES-puro (Halo-βarrestin2) vectors.

### Generation of stable CHO cell lines

Stable low-expression CHO cell lines expressing SNAP_f_-MOR and SNAP_f_-MOR11S/T-A were generated as previously described.^23^ Briefly, we used a Flp-In T-REx (FITR) CHO cell line that had been selected for an FRT recombination target site conferring low expression levels and that also allows for tetracycline-inducible expression, referred to as LEx-FITR cells. These cells constitutively express the Tet repressor under the control of the CMV promoter and maintain the integrated transgene under control of the crippled CMV promoter and two tetracycline operator 2 (2×TetO2) sites. Stable integration of the pcDNA5/FRT/TO-IRES SNAP_f_-MOR vector was achieved by cotransfecting 200 ng of the receptor vector with 1,800 ng of the pOG44 Flp-In recombinase vector (Invitrogen) using Lipofectamine LTX according to the manufacturer’s protocol. Stable integrants were selected with 500 µg/mL hygromycin B (Thermo Fisher Scientific) in the presence of 15 µg/mL blasticidin to maintain both the integrated transgene and the T-REx expression cassette.

### Transient transfections

Transient transfections were performed in six-well format 48 hours before imaging. For HaloTag constructs (β₂AR, V2R, GRK2, and βarrestin2), 250 ng DNA per construct was diluted in 150 µL Opti-MEM with 3 µL Plus reagent (Thermo Fisher Scientific). Separately, 10 µL Lipofectamine LTX (Thermo Fisher Scientific) was diluted in 150 µL Opti-MEM. After 5 min, DNA and lipid mixes were combined and incubated for 20 min before addition to ∼70% confluent CHO cells in 6-well plates containing 2 mL antibiotic-free Ham’s F-12 medium. Following overnight incubation (12–16 h), media were replaced with complete growth medium, and cells were allowed to recover for 24 h prior to labeling or imaging. For experiments requiring tetracycline induction of SNAP_f_ constructs, cells were treated with 0.5 µg/mL tetracycline for 12–16 h before imaging to achieve controlled expression levels suitable for single-molecule analysis.

### Live-cell single-molecule TIRF microscopy and imaging

Single molecules were imaged using a previously described custom-built objective-based TIRF microscope (IX81, Olympus) equipped with a 4 channel fully motorized TIRF illuminator (cell-TIRF, Olympus) and a high numerical aperture oil immersion objective (UAPON 100×/1.49 NA, Olympus).^23,65^ Focus drift was controlled during all measurements by using a laser autofocus system (ZDC2, Olympus).

LD555p-BG was excited with a 532-nm laser (Torus 150 mW, Laser Quantum), and Halo-JF646 was excited with a 640-nm laser (cell laser, 100 mW, Olympus). Excitation and emission light were separated using a dual-band laser filter set (ZT532/640rpc, ZET532/640m, ZET532/640x; Chroma). For dual-color imaging, the fluorescence light was further separated by an OptoSplitter II image splitter (Cairn Research) with a dichroic filter set (ZT640rdc, ET585/65m, ET655lp, Chroma) to project the LD555p and JF646 emission side by side onto an EMCCD camera (Evolve 512, Photometrics). Movies were recorded at 15 ms per frame with an effective pixel size of 160 nm.

#1.5H index glass coverslips were cleaned by base-peroxide bath (H₂O₂:NH₄OH:H₂O, 1:0.25:5 at 75–80 °C, 90 min), manual rinse (dH₂O followed by 100% ethanol), air-drying, and plasma cleaning (5 min; Zepto LF). Coverslips were coated with human plasma fibronectin (10 µg/mL, 30 min), rinsed, assembled in chambers, and equilibrated in imaging buffer before seeding cells.

To maintain temperature stability, cells were imaged at 37 °C in a microscope warming box (Precision Plastics Inc.). For each experimental condition, movies with a length of 4,000 frames were collected, providing sufficient trajectories for diffusion analyses. For low expression conditions, cells were not treated with tetracycline and labeling was performed with LD555p-BG at 500 nM for 15–20 min at 37 °C, followed by five washes with DPBS + 0.1% BSA and recovery in complete medium for 40–60 min. For high expression conditions, cells were treated with 1 µg/mL tetracycline for 12–16 h to achieve higher surface expression. Labeling was performed with 250 pM LD555p-BG to sparsely labeling for suitable particle detection. For dual-color experiments, Halo-tagged constructs were labeled with Halo-JF646 (100 nM) concurrently.

Cells were treated with 1 µM DAMGO (Sigma-Aldrich) for 30 min at 37 °C prior to imaging. For co-stimulation experiments, DAMGO was applied concurrently with quinpirole (10 µM), isoproterenol (10 µM), or AVP (10 µM). For GRK2/3 inhibition, cells were pre-treated with CMPD101 (30 µM, 30 min, 37 °C) followed by DAMGO + CMPD101 for an additional 30 min.

### Surface density determination

Surface density was determined as previously described.^23^ An initial image acquired using simultaneous 532-nm and 640-nm excitation was used to determine the density of fluorescently labeled molecules in the green and red emission channel before single particle tracking. Single molecule counts were determined using the DoG particle-detection function of TrackMate in ImageJ.^66,67^ The cell area was determined by boundary tracing of a projected image, generated from a donor image stack using the ImageJ plugin ZProject with projection type ‘Standard Deviation’.^67^

### Single-particle tracking and motion classification

Single particle tracking (SPT) was performed using u-track, a multi-particle tracking software.^68^ A detailed list of all u-track parameters used in this study can be found in **Table S1**. For subsequent motion analysis of the single-molecule trajectories, Divide-and-Conquer Moment Scaling Spectrum (DC-MSS) was used,^26^ which can directly import u-track results to perform a transient mobility analysis.

### ELISA-based receptor internalization assay

Cells were seeded and transfected two days before experiments, replated the following day, and treated with agonists after allowing 24 hours for recovery. Following treatment, cells were washed with Tris-buffered saline (TBS) and fixed with 4% paraformaldehyde for 30 minutes at room temperature. Fixed cells were blocked with 0.1 M sodium bicarbonate (pH 8.6) containing 1% milk for 4 hours at room temperature or overnight at 4 °C. After blocking, cells were incubated with anti-HA primary antibody (1:1000, TBS + 0.1% BSA (Sigma-Aldrich)) to label surface receptors, followed by horseradish peroxidase (HRP)–conjugated anti-mouse secondary antibody (1:10,000 (Sigma-Aldrich)) in fresh blocking buffer.

After final washes with TBS, colorimetric detection was performed using OPD substrate (Thermo Fisher). The reaction was quenched with 1 M sulfuric acid, and absorbance was measured at 490 nm (A490). Signals from mock-transfected or uninduced wells were used to define the background and subtracted from all readings. Absorbance at 490 nm served as a measure of surface receptor abundance. Internalization was quantified as the fractional loss of surface receptor signal relative to untreated controls.

### TIRF-based colocalization microscopy

For TIRF clathrin colocalization, imaging was performed on an Olympus IX83 with CellTIRF1 illuminator, UApoplan 60×/1.49 NA TIRF objective, Toptica iChrome MLE laser engine (488 nm and 561 nm, 100 mW each), Andor Zyla 4.2 sCMOS camera, and a Chroma TRF89901-OL3 quad-band filter cube. Emitted fluorescence was further purified using filter sets equipped with 495LP dichroic beamsplitter and 540/25 emission filter for the 488nm laser line, and 565LP dichroic beamsplitter and 605/50 emission filter for the 561nm laser line (Chroma). Temperature was maintained at 37 °C using objective and stage heaters (Bioptechs Stable Z). Single-molecule images were acquired at 50 ms exposure using the 561 nm laser (25 mW). CLC-mNeonGreen images were acquired using 1000 ms exposure. Lateral shift between the different detection channels was determined using Tetraspeck 40 nm beads as a control.

### Image registration, detection, and colocalization analysis

TIRF colocalization analysis has been described previously.^69^ Particle detection for single molecules employed the Fiji TrackMate LoG detector with detection diameter of 0.4 µm, median filter, standardized quality thresholding, and subpixel localization precision.^66,70^ Localization of clathrin-coated pits was done using adaptive local thresholding based on Niblack’s algorithm with fixed parameters.The drift correction between individual channels was performed using NanoJ-Core Fiji plugin using Tetraspeck bead data as a reference.^71^ For colocalization estimation positions of detected molecules were matched with coordinates of clathrin-coated pits and the fraction of colocalized molecules were obtained. Fraction measurements were subsequently adjusted for possible random colocalizations to determine a colocalization coefficient which was used to calculate the fraction of non-randomly colocalized molecules.

### cAMP assay

The bioluminescence resonance energy transfer (BRET)-based cAMP inhibition assay was carried out in the SNAP_f_-MOR stable LEx-FITR CHO line. For the MOR-mediated cAMP inhibition assay, the plasmid encoding the cAMP sensor using YFP-Epac-RLuc (CAMYEL, 1 μg, ATCC)^72^ was transfected. Cells were prepared and assayed as described previously in detail using a 96-well microplate and PHERAstar FS plate reader.^23^ After incubation with 10 μM forskolin (Cayman Chemical Company) and incubation with coelenterazine h (5 µM, NanoLight Technology) for 7 minutes, [D-Ala2, NMePhe4, Gly-ol]-enkephalin (DAMGO) (MilliporeSigma) was injected. BRET measurements were taken 20 minutes after injection of ligand. Data analysis was carried out as described previously^23^.

### Plotting and statistics

Plotting, distribution fitting and statistics for all single-molecule and colocalization data were carried out using GraphPad Prism (GraphPad Software). To determine *P* values, two-way ANOVA with Tukey’s post hoc test was performed for multisample comparisons, while an unpaired two-sided *t*-test was used for two-sample comparisons.

